# Biasing AlphaFold2 to predict GPCRs and Kinases with user-defined functional or structural properties

**DOI:** 10.1101/2022.12.11.519936

**Authors:** D. Sala, J. Meiler

## Abstract

Determining the three-dimensional structure of proteins in their native functional states has been a longstanding challenge in structural biology. While experimental methods combined with integrative structural biology has been the most effective way to get high accuracy structures and mechanistic insights for larger proteins, advances in deep machine-learning algorithms have paved the way to fully computational predictions. In this field, AlphaFold2 (AF2) pioneered *ab initio* high accuracy single chain modeling. Since then, different customizations expanded the number of conformational states accessible through AF2. Here, we further extended AF2 with the aim of enriching an ensemble of models with user-defined functional or structural features. We tackled two common protein families for drug discovery, G-protein-coupled receptors (GPCRs) and Kinases. Our approach automatically identifies the best templates satisfying the specified features and combines those with genetic information. We also introduced the possibility of shuffling the selected templates to expand the space of solutions. In our benchmark, models showed the intended bias and great accuracy. Our protocol can thus be exploited for modeling user-defined conformational states in automatic fashion.

## INTRODUCTION

X-ray crystallography and cryogenic electron microscopy (cryo-EM) are two widely used techniques for determining the detailed structures of biomolecules at the atomic level [1,2]. For structure-based drug discovery and design, having at least one high accuracy structure is essential [3]. Despite recent advances in technology that have made more protein structures available [4], their experimental determination is still a difficult and costly process with a high risk of failure [5]. In fact, experimental protein structures represent only a small fraction of the complete set of known protein sequences [6,7]. Furthermore, one structure only represents a snapshot of a protein, and may not necessarily be sufficient to understand the overall mechanism of operation. This limitation has important implications for drug discovery, especially for common drug targets such as G-protein-coupled receptors (GPCRs) and kinases, which are known to modulate cellular behavior by switching between multiple structurally different functional states [8,9].

The 14^th^ edition of Critical Assessment of protein Structure Prediction (CASP14) has recognized AlphaFold2 (AF2) for its impressive accuracy in predicting monomeric protein structures *de novo* [10]. AF2 makes it straight-forward to predict a protein structure from a protein sequence, and has provided millions of protein models with estimated accuracy [11]. Since the emergence of AF2, a number of deep learning-based methods have been developed with the same goal of predicting protein structures at experimental accuracy [12–15]. Among them, RoseTTAFold was the first approach that was able to predict both active and inactive GPCR conformations, outperforming comparative homology modeling by using templates in a uniform functional state [12]. This achievement has sparked interest in developing workflows to predict multiple native conformations of a protein target with the state-of-the-art AF2 implementation.

To date, a number of AF2 customizations that adopted different concepts are available [16–19]. Del Alamo and co-authors took advantage of a shallow multiple sequence alignment (sMSA) to collect an ensemble of conformations, among which multiple native conformations of GPCRs and transporters were identified [16]. Alternatively, SPEACH_AF (hereafter SPEACH) masked multiple positions in the multiple sequence alignment (MSA) to switch the prediction toward alternative conformational states that were less represented in the MSA [18]. Another protocol removed the MSA (noMSA) and prepared a local database of state-annotated GPCRs to perform AF2 template-based modelling [17]. All methods for sampling conformational changes in proteins have shown great potential, but also have some limitations, such as a reduced breadth of sampled conformations or a high dependence on the structural features of selected templates.

Here, we update our previous protocol (sMSA) to facilitate the collection of templates with user-defined functional or structural properties of GPCRs and kinases. Templates are automatically filtered and retrieved from an annotated database in accord to the specified functional or structural criteria. Through a calibrated balancing of genetic and template-based features, our protocol samples equal or better active GPCR models than all the released methods for sampling alternative states. On a difficult target, randomizing templates to explore the available structural space significantly improved accuracy. In modeling kinase conformations, our protocol enriched the predicted ensemble with models carrying user-defined structural features.

## METHODS

We updated our previous modified ColabFold version [16,20] and our python interface to allow users to specify functional or structural properties of templates for modeling GPCRs and kinases. The new implementation and accompanying documentation with examples can be found at https://github.com/davidesala/af2_conformations.

### GPCRs benchmark

Target PDBs for LSHR, MTR1A, PE2R4 and ADRB1 were 7FII, 7VGY, 7D7M and 7JJ0, respectively [21–24]. The protein regions corresponding to transmembrane helices (TM-RMSD) were retrieved from GPCRdb [25]. Four workflows were evaluated to predict the alternative conformational states of GPCRs: ActTemp+sMSA was run with 8 sequence clusters and 16 extra clusters sequences combined with the automatic detection of the top 4 ‘Active’ templates. For LSHR, MTR1A, PE2R4 and ADRB1 those were: 6H7L_A-6IBL_A-6K41_R-6K42_R, 6H7L_A-7P00_R-6IBL_A-7RMG_R, 7E32_R-7CKY_R-7CKW_R-7JVP_R and 6MXT_A-7CKY_R-7CKW_R-7JVP_R, respectively. Other AF2 parameters were kept as in our previous pipeline - named sMSA - that used 16 sequence clusters and 32 extra clusters sequence without any template and no recycling [16]. To remove the MSA (noMSA run), the same implementation published previously was adopted [17]. These runs were then carried out using the GPCRdb API rather than a local GPCR database to avoid mismatches between the pool of available templates. The SPEACH protocol was applied with a sliding window of 10 masked residues [18]. Thus, the number of models collected with SPEACH was higher than the 50 models collected with other protocols. Models with a pTM score < 70 were found to be unfolded and discharged.

To assess the impact of randomizing templates, the inactive state structure of LT4R1 (PDB 7K15) was used as a target [26]. The MSA for the aligned regions was removed, and 50 models were generated with and without randomizing templates. The templates used for the models without randomization were 6VI4_A-4ZUD_A-4YAY_A-4N6H_A.

### EIF2AK4 kinase benchmark

Model were predicted by using the same parameters of the new ActTemp+sMSA implementation but with 20 templates instead of 4. The DFG, aC_helix and Salt bridge K^III.17^ and E^αC.24^ structural features as well as the activation loop orientation were defined according to the KLIF database [28]. Unfolded models were discarded.

## RESULTS

The original pipeline that was developed to sample alternative conformations was expanded to improve the prediction of GPCRs and kinases in a specific conformational state. Users can now specify the activation state of GPCRs and the script will look for templates that match that state or are bound to a signaling protein. To do so, one of the following labels must be declared: ‘Active’, ‘Inactive’, ‘Intermediate’, ‘G protein’, ‘Arrestin’. For kinases, users can select specific structural features values and the script will search for templates that match those criteria. Allowed values for the corresponding structural feature are: i) ‘out’, ‘in’, ‘out-like’, ‘all’ for DFG, ii) ‘out’, ‘in’, ‘all’ for aC_helix, iii) ‘yes’, ‘no’, ‘all’ for Salt bridge K^III.17^ E^αC.24^ [27]. Optionally, the list of templates that pass the sequence and structural filters can be randomized to explore the available structural space.

In the sections below, we demonstrate how selecting templates in accord to functional or structural properties and combining those with genetic information can influence the predicted structural features of the models. We also show the results of randomizing templates on a difficult target.

### Combining a shallow MSA with state-annotated templates outperforms state-of-the-art methods for sampling GPCRs active state

Our new pipeline was used to predict GPCR models by combining a very shallow MSA with the automatic detection of the best 4 active templates from GPCRdb (ActTemp+sMSA). The benchmark set of these GPCRs consisted of four proteins: LSHR, MTR1A, PE2R4 and ADRB1. The first three were previously identified as challenging to predict in the active state [17]. Instead, ADRB1 was solved in the inactive conformation before April 2018 and therefore was part of the AF2 training set. For this reason, we included in the benchmark set the more recent ADRB1 active state structure with the specific aim of assessing methods ability to overcome the AF2 inactive state bias. For each method, we measured the accuracy as Cα-RMSD (root-mean-square deviation) of the transmembrane helices (TM-RMSD) with respect to the experimental structure. Our implementation was compared to all the AF2 workflows designed to sample alternative protein conformations. ActTemp+sMSA was the only approach that consistently generated models with near or subangstrom accuracy for all the GPCRs (Figure 1). Interestingly, our approach and noMSA were the only methods able to overcome the ADRB1 inactive state bias and accurately model the active state with an average accuracy of 0.5 Å.

**Figure 1.**
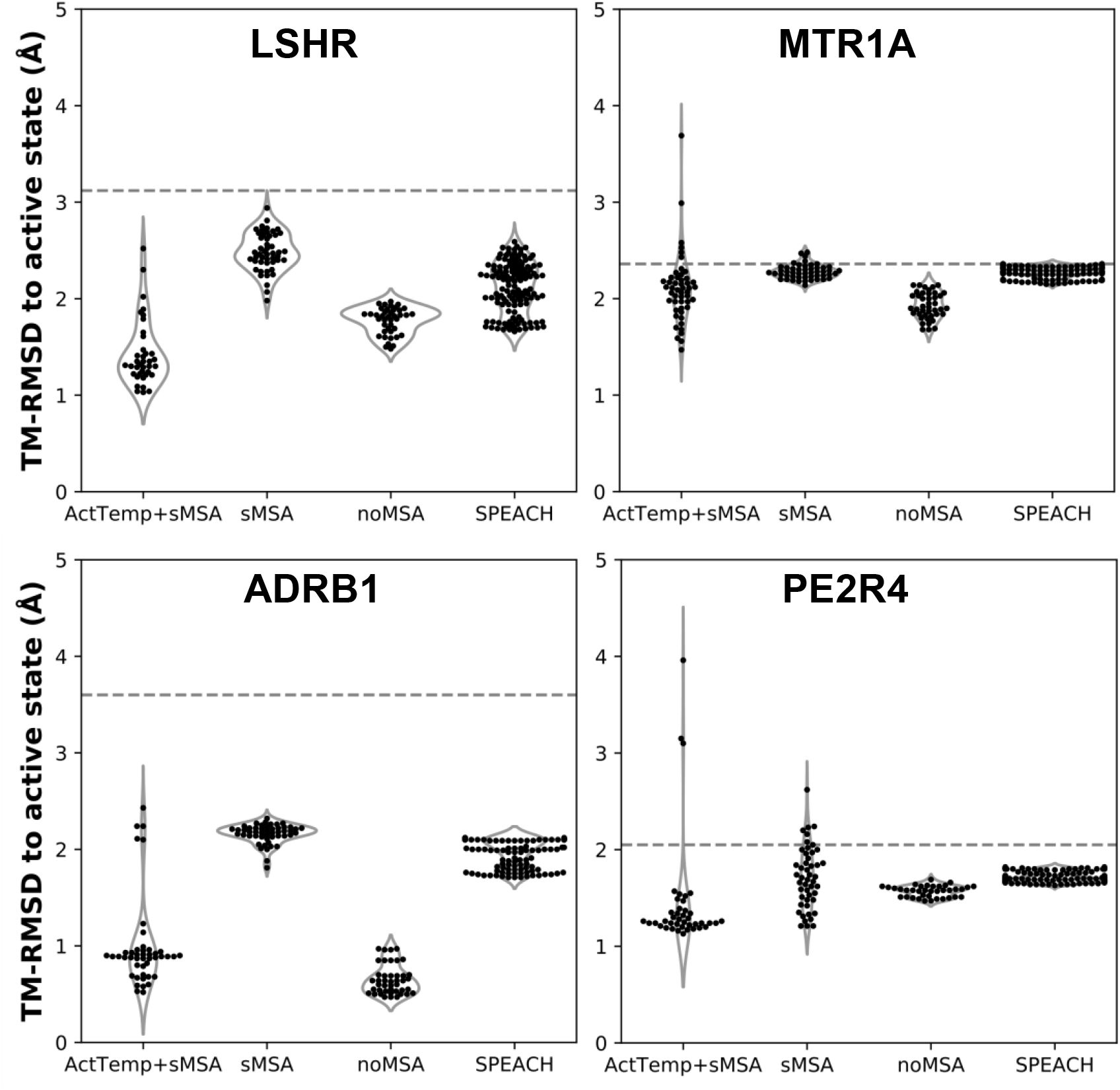
AF2 accuracy in predicting active state GPCRs with different protocols. ActTemp+sMSA was predicted with templates in the active state and a shallow MSA, sMSA with a shallow MSA only, noMSA without a MSA for templates aligned regions, SPEACH with a sliding window masked MSA. TM-RMSD between experimental active and inactive structures is shown as a dashed line.

### Shuffling templates in a homogenous functional state can improve accuracy

Given that subsampling the sequence space (i.e., the MSA) returns different models, we hypothesized that randomly selecting a subset of templates can potentially yield more accurate models. To test this, we removed the genetic information within the AF2 pipeline and generated 50 models with and without randomizing inactive templates. For each model, our script selected 4 random inactive state structures from GPCRdb that passed the sequence similarity filter. Accuracy was measured as TM-RMSD from the inactive state structure of LT4R1 (PDB 7K15). The exploration of the structural space defined by the ensemble of all the inactive templates resulted in more accurate models compared to using the top 4 templates (Figure 2A).

**Figure 2.**
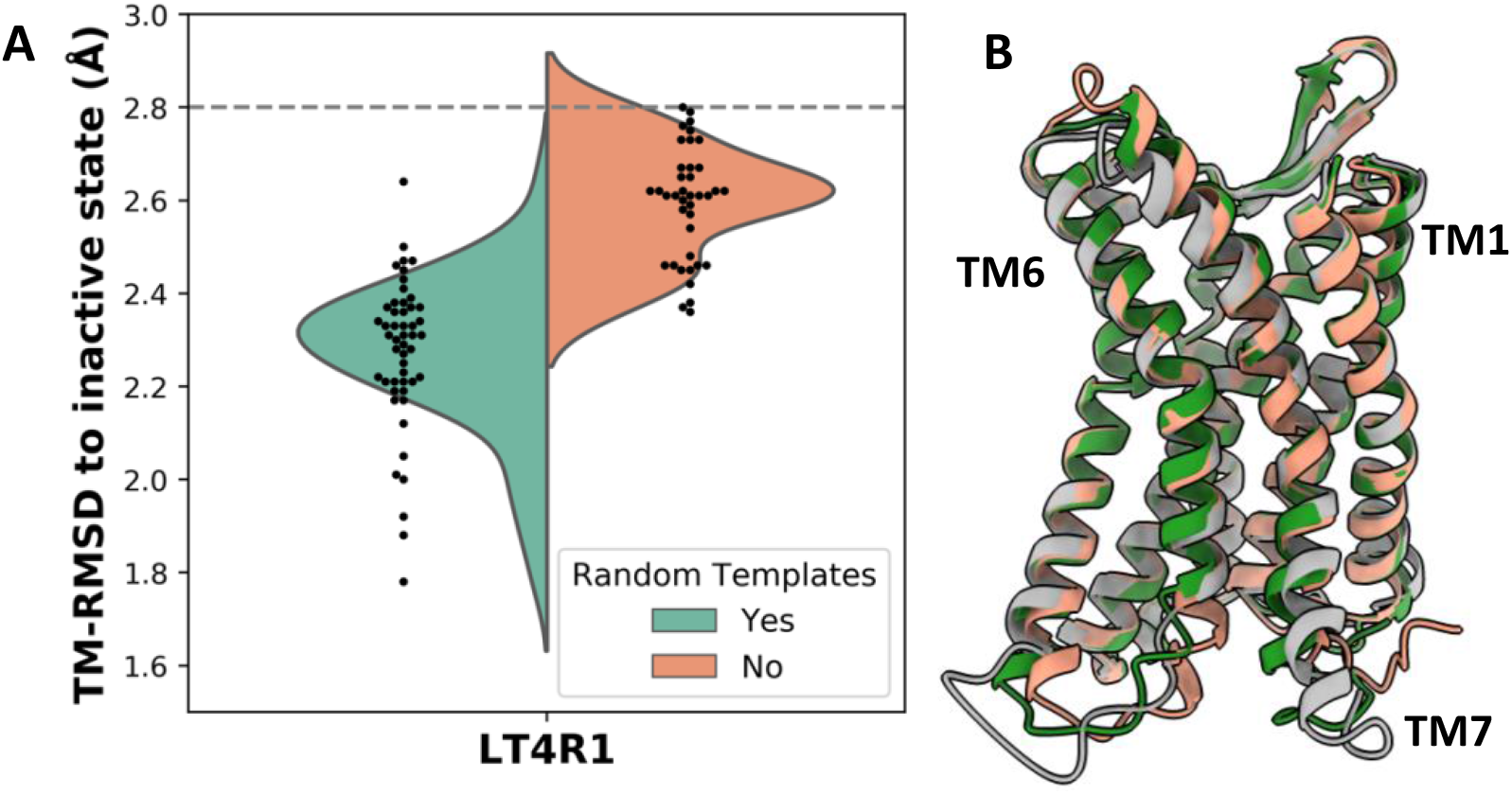
Accuracy in predicting the LT4R1 inactive state with and without randomizing templates. A) TM-RMSD distribution of models. TM-RMSD between experimental active and inactive structures is shown as a dashed line. B) Superposition of the best model from the random templates ensemble (green) and without randomizing templates (orange) to the experimental structure (gray).

The superposition of the best model in the two ensembles shows improved fitting of the long TM7 helix and better modeling of TM1 and TM6 when using random templates (Figure 2B).

### User-defined structural features to bias kinase modeling

The concept of allowing users to define structural features of GPCR templates was also applied to kinases using the KLIF webserver. We implemented the possibility to choose three conformational properties: DFG, αC-helix (ac_H) and salt bridge K^III.17^E^αC.24^. The script automatically selects and retrieves templates satisfying user-defined structural criteria. We assessed the effect on the predicted conformations by modeling the EIF2AK4 (GCN2) kinase. We generated 50 models for each of the following templates biased features: ‘DFG=all/ac_H=all’ which is equivalent to predict without any bias; ‘DFG=in/ac_H=in’ and ‘DFG=in/ac_H=out’, which differ in the αC-helix position regardless of its rotation; and ‘DFG=out/ac_H=all’, which corresponds to the inactive state required for binding type II inhibitors. Because DFG is a multi-criteria parameter, we also evaluated the predicted position of the activation loop (a_loop), which is well-defined and mostly corresponds to DFG. Without biasing the prediction, most of the models were found in the ‘a_loop=in/ac_H=out’ conformation, while 20% of the pool was in the ‘a_loop=in/ac_H=in’ conformation, and only one model was found with ‘a_loop=out’ (Figure 3). By keeping ‘DFG=in’ while changing ac_H templates, AF2 generated most of the models in agreement with the corresponding templates ac_H position. The bias was slightly stronger for ‘ac_H=out’ than ‘ac_H=in’, following the same trend as the prediction without any bias. While ‘DFG=in’ templates generated only ‘a_loop=in’ conformations, ‘DFG=out’ templates significantly increased the number of models in the inactive state carrying the crucial ‘a_loop=out’ conformation (green bar).

**Figure 3.**
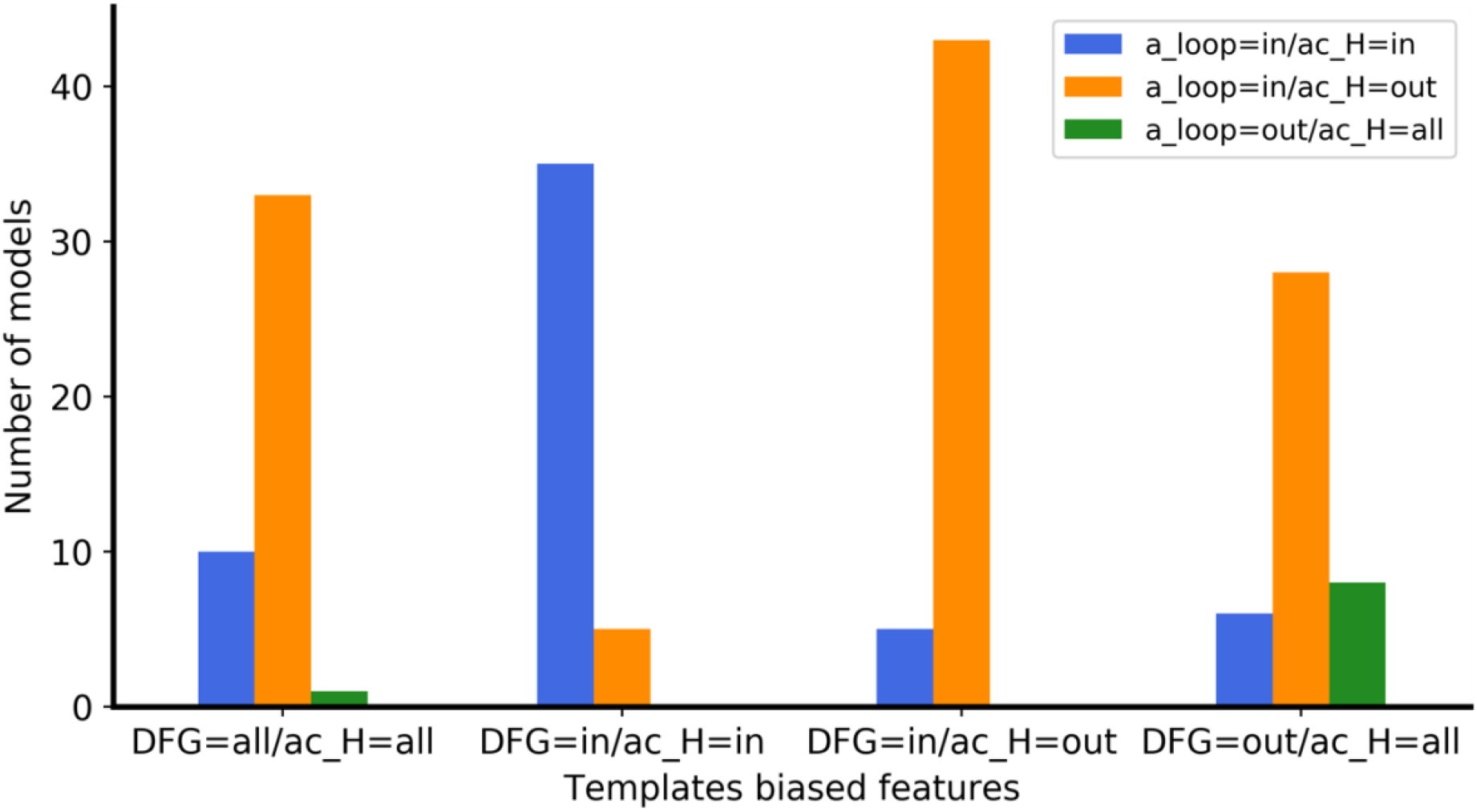
Enrichment of eif2k4 kinase models with structural properties corresponding to the biased templates features used. The four ensembles were calculated with a different ‘DFG/ac_H’ templates bias. For each ensemble, the number of models with the three ‘a_loop/ac_H’ conformational feature combinations is shown with a different color bar.

The superimposition of ‘a_loop=out’ and ‘a_loop=in’ models onto the experimental ‘DFG=out’ (PDB 7QWK) and ‘DFG=in’ structures (PDB 7QQ6) shows that they fit the experimental DFG loops orientation, with a small discrepancy for ‘DFG=out’ likely due to the presence of the inhibitor in the experimental structure (Figure 4) [29].

**Figure 4.**
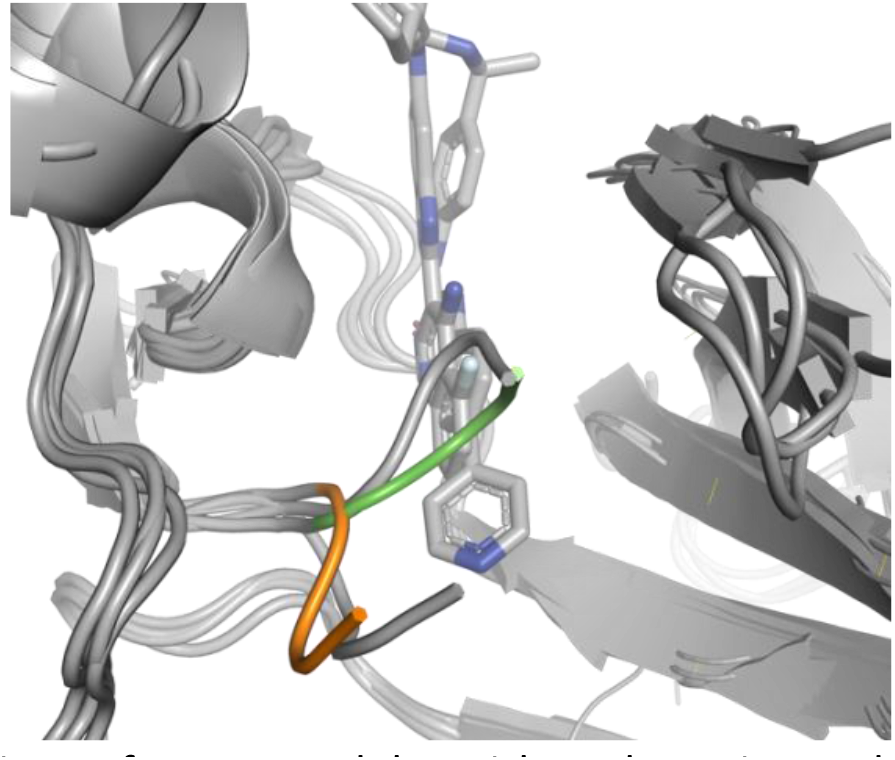
Superposition of two models with a_loop=in and a_loop=out to the two corresponding ‘DFG=in’ and ‘DFG=out’ experimental structures. DFG residues of models with ‘out’ and ‘in’ orientations are shown in green and orange, respectively. Experimental structures of eif2k4 are shown in gray.

## DISCUSSION

The prediction of user-defined conformational states of proteins has been a challenge even after the advent of AF2. Previous workflows attempting to solve this problem either do not explicitly predict user-defined structural properties or require the creation of state-annotated local structure databases [16–19]. In this work, we propose a pipeline that biases AF2 predictions toward the intended functional state of GPCRs or toward specific structural properties of kinases. One key aspect of our method is its simplicity in use. By leveraging on the APIs of two popular webservers, GPCRdb and KLIFS, our script filters templates according to pre-defined structural or functional parameters, allowing for fully automatic selection of templates without the need for manual inspection or for downloading and updating of databases.

Our results in predicting the active structures of several challenging GPCRs show that combining a shallow multiple sequence alignment (MSA) with templates in a user-defined activation state outperforms existing AF2 workflows. Our pipeline generated models with near or subangstrom accuracy. A direct comparison with models predicted without an MSA (noMSA) suggests that the balanced combination of genetic (MSA) and structural (templates) features may be crucial for achieving high accuracy. This balanced mixture enables a structural refinement while avoids the overwhelming effect coming from a deep MSA, as previously reported [16]. Another advantage of a balanced mixture of genetic and structural information is its reduced sensitivity to neural network biases. In our benchmark, target conformations were all class A GPCRs for which inactive structures were more prevalent than active ones in the AF2 training set. Furthermore, the inactive structure of ADRB1 was directly part of the AF2 training set, thus representing a very strong bias. Indeed, protocols relying solely on genetic information (sMSA and SPEACH) were on average less accurate and completely missed the target conformation for ADRB1.

Our efforts to bias the prediction of a kinase toward user-defined structural properties exploited two important structural components that define its activation state: DFG and αC-helix. While the latter was easier to direct toward the intended position, the former was more difficult likely due to the neural network bias in the training set composition. Despite this, we successfully generated multiple models with ‘DFG=out’ conformation. Given that ‘DFG=out’ structures are needed for structure-based drug design and discovery of type-II inhibitors [30], our script is well positioned to generate models carrying this crucial structural feature.

Our work expands the portfolio of AlphaFold2 customizations developed with the aim of predicting multiple conformational states of proteins. Our python interface facilitates the prediction of intended functional or structural properties of GPCRs and kinases and can be further extended to include more properties as needed. We also emphasize the importance that structure- and function-annotated databases had for this work. The expansion of existing databases to include additional annotations and the development of new protein family-based databases would improve or enable automatic calibrated modeling, respectively. Together, curated databases and machine learning offer a powerful combination for high throughput modeling and, ultimately, for structure-based drug discovery.

## ACKNOWLEDGEMENTS

This work was supported by the Deutsche Forschungsgemeinschaft (DFG, German Research Foundation) through CRC 1423, project number 421152132, subproject A07 and Z04. Jens Meiler is supported by a Humboldt Professorship of the Alexander von Humboldt Foundation. We thank the Deutsche Forschungsgemeinschaft (SPP 2363, “Molecular Machine Learning”) for generous financial support. The work was further supported by NIH NIGMS R01 GM080403, NIH NIHL R01 HL122010, and NIH NIDA R01 DA046138. The authors would like to thank Dr. Ben Brown for useful discussions.

## DECLARATION OF INTERESTS

No interests are declared.

## Notes

### Competing Interest Statement

The authors have declared no competing interest.

## REFERENCES

1 Wang, H.W. and Wang, J.W. (2017) How cryo-electron microscopy and X-ray crystallography complement each other. Protein Sci. 26, 32–39

2 Vénien-Bryan, C. et al. (2017) Cryo-electron microscopy and X-ray crystallography: Complementary approaches to structural biology and drug discovery. Acta Crystallogr. Sect. Struct. Biol. Commun. 73, 174–183

3 Congreve, M. et al. (2020) Impact of GPCR Structures on Drug Discovery. Cell 181, 81–91

4 Callaway, E. (2020) Revolutionary cryo-EM is taking over structural biology. Nature 578, 201

5 Lyumkis, D. (2019) Challenges and opportunities in cryo-EM single-particle analysis. J. Biol. Chem. 294, 5181–5197

6 The Uniprot Consortium (2019) UniProt: a worldwide hub of protein knowledge. Nucleic Acids Res. 47, D506–D515

7 Burley, S.K. et al. (2021) RCSB Protein Data Bank: powerful new tools for exploring 3D structures of biological macromolecules for basic and applied research and education in fundamental biology, biomedicine, biotechnology, bioengineering and energy sciences. Nucleic Acids Res. 49, D437–D451

8 Attwood, M.M. et al. (2021) Trends in kinase drug discovery: targets, indications and inhibitor design. Nat. Rev. Drug Discov. 20, 839–861

9 Yang, D. et al. G protein-coupled receptors: structure-and function-based drug discovery., Signal Transduction and Targeted Therapy, 6. 08-Jan-(2021), Nature Publishing Group, 1–27

10 Jumper, J. et al. (2021) Highly accurate protein structure prediction with AlphaFold. Nature 596, 583–589

11 Tunyasuvunakool, K. et al. (2021) Highly accurate protein structure prediction for the human proteome. Nature 596, 590–596

12 Baek, M. et al. (2021) Accurate prediction of protein structures and interactions using a three-track neural network. Science (80-.). 373, 871–876

13 AlQuraishi, M. Machine learning in protein structure prediction., Current Opinion in Chemical Biology, 65. 01-Dec-(2021), Elsevier Current Trends, 1–8

14 Chowdhury, R. et al. (2022) Single-sequence protein structure prediction using a language model and deep learning. Nat. Biotechnol. 40, 1617–1623

15 Lin, Z. et al. (2022) Evolutionary-scale prediction of atomic level protein structure with a language model. bioRxiv DOI: 10.1101/2022.07.20.500902

16 del Alamo, D. et al. (2022) Sampling alternative conformational states of transporters and receptors with AlphaFold2. Elife 11, 1–12

17 Heo, L. and Feig, M. (2022) Multi-state modeling of G-protein coupled receptors at experimental accuracy. Proteins Struct. Funct. Bioinforma. DOI: 10.1002/prot.26382

18 Stein, R.A. et al. (2022) SPEACH_AF: Sampling protein ensembles and conformational heterogeneity with Alphafold2. PLOS Comput. Biol. 18, e1010483

19 Wayment-Steele, H.K. et al. Prediction of multiple conformational states by combining sequence clustering with AlphaFold2. DOI: 10.1101/2022.10.17.512570

20 Mirdita, M. et al. (2022) ColabFold: making protein folding accessible to all. Nat. Methods 19, 679–682

21 Su, M. et al. (2020) Structural Basis of the Activation of Heterotrimeric Gs-Protein by Isoproterenol-Bound &#x3b2;_1_-Adrenergic Receptor. Mol. Cell 80, 59–71.e4

22 Duan, J. et al. (2021) Structures of full-length glycoprotein hormone receptor signalling complexes. Nature 598, 688–692

23 Nojima, S. et al. (2021) Cryo-EM Structure of the Prostaglandin E Receptor EP4 Coupled to G Protein. Structure 29, 252–260.e6

24 Wang, Q. et al. (2022) Structural basis of the ligand binding and signaling mechanism of melatonin receptors. Nat. Commun. 13, 454

25 Kooistra, A.J. et al. (2021) GPCRdb in 2021: Integrating GPCR sequence, structure and function. Nucleic Acids Res. 49, D335–D343

26 Michaelian, N. et al. (2021) Structural insights on ligand recognition at the human leukotriene B4 receptor 1. Nat. Commun. 12, 2971

27 McClendon, C.L. et al. (2014) Dynamic architecture of a protein kinase. Proc. Natl. Acad. Sci. 111, E4623–E4631

28 Kanev, G.K. et al. (2021) KLIFS: an overhaul after the first 5 years of supporting kinase research. Nucleic Acids Res. 49, D562–D569

29 Maia de Oliveira, T. et al. (2020) The structure of human GCN2 reveals a parallel, back-to-back kinase dimer with a plastic DFG activation loop motif. Biochem. J. 477, 275–284

30 Ung, P.M.-U. and Schlessinger, A. (2015) DFGmodel: Predicting Protein Kinase Structures in Inactive States for Structure-Based Discovery of Type-II Inhibitors. ACS Chem. Biol. 10, 269–278

